# Slow waves form expanding, memory-rich mesostates steered by local excitability in fading anesthesia

**DOI:** 10.1101/2021.01.21.427671

**Authors:** Antonio Pazienti, Andrea Galluzzi, Miguel Dasilva, Maria V. Sanchez-Vives, Maurizio Mattia

**Affiliations:** Natl. Centre for Radioprotection and Computational Physics, Istituto Superiore di Sanità, Rome, Italy; IDIBAPS (Institut d’Investigacions Biomèdiques August Pi i Sunyer), Barcelona, Spain; ICREA (Institució Catalana de Recerca i Estudis Avançats), Barcelona, Spain

## Abstract

During sleep and anesthesia, large groups of neurons throughout the entire cortex activate rhythmically producing wavefronts of activity referred to as slow-wave activity (SWA). In the arousal process, the brain restores its integrative and complex activity. The network mechanisms underlying this global state transition remain however to be elucidated. Here we investigated the network features shaping the SWA under fading anesthesia. Using electrocorticographical recordings of wide cortical areas of the mouse brain, we developed a quantitative measure of the anesthesia level based on slow-wave frequency and complexity. At deep anesthesia, we document a stringent alternation of posterior-anterior-posterior modes of propagation. With fading anesthesia, SWA evolves to produce a plethora of metastable spatiotemporal patterns. A network model of spiking neurons reproduced the data using short-range connectivity, subcortical input and a local activity-dependent adaptation. The emergence from deep anesthesia does not require modifying the connectivity, but small changes in the local excitability of cortical cell assemblies, further supporting the hypothesis of a tight bound between scales in the brain.

## Introduction

During sleep and anesthesia, the brain spontaneously generates patterns of low-frequency, synchronized activity referred to as slow-wave activity (SWA) (Destexhe et al., 1999; Massimini et al., 2004; Mohajerani et al., 2010; Steriade et al., 1993b). This activity coordinates large portions of the cortex into a coherent rhythmic sequence of recurring cortical and cortico-thalamic activations (Destexhe and Sejnowski, 2003; Grenier et al., 1998; Sheroziya and Timofeev, 2014). In this state, cortical neurons undergo a slow alternation between periods of depolarization and firing (Up or On states) and periods of almost-silent hyperpolarization (Down or Off states) (Steriade et al., 1993c; Torao-Angosto et al., 2021; Vyazovskiy et al., 2009). This is accompanied by reduced amounts of neuronal excitability and enhanced segregation of cortical areas (Bettinardi et al., 2015; Compte et al., 2003; Fernandez et al., 2017; Fischer et al., 2018; Levenstein et al., 2019; Liu et al., 2013; Massimini et al., 2005, 2004; Mattia and Sanchez-Vives, 2012; Muller and Destexhe, 2012; Sanchez-Vives and Mattia, 2014; Sanchez-Vives and McCormick, 2000).

In the last two decades, several studies have performed recordings of the neuronal activity during various depths of either natural sleep or anesthesia, as well as during the process of awakening (Barttfeld et al., 2015; Bettinardi et al., 2015; Dasilva et al., 2021; Fischer et al., 2018; Hahn et al., 2012; Hudetz et al., 2015; Hudson et al., 2014; Lee et al., 2020; Li and Mashour, 2019; Liu et al., 2013; Schartner et al., 2017; Tort-Colet et al., 2019). Results from single – or a small number of simultaneously recorded – areas (including primary visual (Hudetz et al., 2015; Lee et al., 2020; Vizuete et al., 2012), cingulate (Hudson et al., 2014), retrospenial (Hudson et al., 2014), temporo-parieto-occipital (Fischer et al., 2018) cortices and the thalamus (Hudson et al., 2014)), as well as from the whole cortex (Barttfeld et al., 2015; Bettinardi et al., 2015; Dasilva et al., 2021; Grandjean et al., 2014; Li and Mashour, 2019; Liu et al., 2013; Schartner et al., 2017), showed that during the emergence from deep anesthesia (NREM sleep) the network activity increases its integration and complexity dynamical properties, possibly starting to wander among so-called micro-states (Brodbeck et al., 2012; Hudson et al., 2014; Lee et al., 2020; Liu et al., 2013). These studies strengthened the hypothesis that the anesthetized (sleeping) brain is not confined in a static dynamical state, but actually explores a more complex landscape, eventually operating a rather progressive state transition to wakefulness (Barttfeld et al., 2015; Dasilva et al., 2021; Li and Mashour, 2019; Stevner et al., 2019).

Several mechanisms are thought to play a role during the emergence of slow-wave states, and during the inverse process that leads to awakening (Deco et al., 2014; Destexhe and Sejnowski, 2003; Längkvist et al., 2012; Mattia and Sanchez-Vives, 2012; Pearlmutter and Houghton, 2009; Sanchez-Vives et al., 2017; Stevner et al., 2019; Tort-Colet et al., 2019). These include cortico-thalamic loops (Crunelli et al., 2015; Crunelli and Hughes, 2010; Destexhe and Sejnowski, 2003; Grenier et al., 1998; Merica and Fortune, 2004; Sheroziya and Timofeev, 2014; Steriade et al., 1993b) and the balance between local dynamical features and global connectivity of the brain network shaping its integration and segregation capabilities (Barttfeld et al., 2015; Deco et al., 2015, 2014; Mohajerani et al., 2013). At the cellular and local cortical assembly level, various experimental and modelling studies pointed at two main players playing a critical role in the process of awakening: firing adaptation of excitatory cortical neurons, and neuronal excitability (Compte et al., 2003; Lee et al., 2020; Levenstein et al., 2019; Mattia and Sanchez-Vives, 2012; Muller and Destexhe, 2012; Sanchez-Vives and McCormick, 2000). However, the cellular and network mechanisms underlying this state transitions remain to be conclusively elucidated. In particular, an open question is the role played by the restored levels of local excitability and long-range connectivity in shaping the activity observed at different levels of anesthesia as the awake state is approached.

Here, by probing the electrophysiological activity of neuronal populations across a wide area of the mouse cortical surface in the arousal from deep isoflurane anesthesia, we addressed the following questions: what is the functional role and the nature of the micro-states composing SWA? What are the relative roles of local and global network features in shaping the SWA as the anesthesia fades out? To this purpose we developed an objective and quantitative measure of the level of anesthesia based on the slow-wave frequency and the complexity of the wavefronts, referred to as the anesthesia level index (ALI) further advancing our previous characterization (Dasilva et al., 2021). This allowed to sort the recordings from different mice and at varying isoflurane doses on a common fine grained frame. As result, we found a significant correlation between ALI and the number of spontaneously expressed modes of propagation, and the progressive loss and recovery of sequential memory between consecutive waves. Resorting to a spatially-extended spiking neuron network modelling the cortical surface probed in experiments, we reproduced the large majority of the observations found at different levels of anesthesia. The rise of complexity in the model was obtained without modifying the underlying “structure”, i.e. the cortico-cortical connectivity. The changes in the local excitability of model neurons alone were enough to explain the dynamical transition of the global cortical network in the arousal process from deep anesthesia, supporting the tight binding between scales in the brain.

## Results

### Lightening up of anesthesia increases frequency and complexity of slow waves

In order to understand the local and global changes underpinning the progressive transition of the cortical network in the arousal process from deep anesthesia, we recorded the ongoing neuronal activity across three levels of anesthesia (isoflurane) in the intact mouse brain. We placed a multi-electrode array (MEA) of 32 channels on the cortical surface of *n* = 8 wild-type mice (Fig. 1A, left; see Materials and Methods). The probed surface spanned several cortical areas including both sensory and motor cortices of a single hemisphere (Fig. 1A, right). Anesthesia levels were finely tuned to induce slow-wave activity (SWA), i.e. the quasi-periodic alternation between slowly propagating onsets of high-firing Up states and almost quiescent Down periods across the probed surface ((Liu et al., 2013; Ruiz-Mejias et al., 2011; Torao-Angosto et al., 2021; Vijayan et al., 2010); Fig. 1B). Spontaneously occurring spatiotemporal patterns were well visible in the multi-unit activity (MUA) signals although electrophysiological recordings were epicortical and not intraparenchymal (Fig. 1C). This allowed us to characterize the changes in the onsets and offsets of the Up states with the lightening up of the anesthesia levels at multiple spatial and temporal scales. The timing of Down-Up transitions for every channel led to a time-lag matrix (TLM) (Capone et al., 2019; Ruiz-Mejias et al., 2011), where each row corresponds to a wavefront containing the relative delays in the channels activation (Fig. 1B-bottom, Materials and Methods). Computing the principal components (Jolliffe and Cadima, 2016) of the TLM and transforming the eigenvalue distribution of the covariance matrix into probabilities, we estimated the number of the effective dimensions (i.e., the principal components) of the space embedding the activation wavefronts. The number of effective dimensions strongly correlates with the slow-wave frequency across experiments and anesthesia levels (Fig. 1D, *n* = 24). Interestingly, the projection of each of these points onto the first principal component of the distribution (blue line in Fig. 1D) is not only coherent with the corresponding isoflurane level (Fig. 1E), but it also allows to finely classify the experiments according to their effective level of anesthesia. We called the projected value on this axis the “anesthesia level index” (ALI), and used it to visualize the progression of other relevant quantities across the different anesthesia levels.

**Figure 1:**
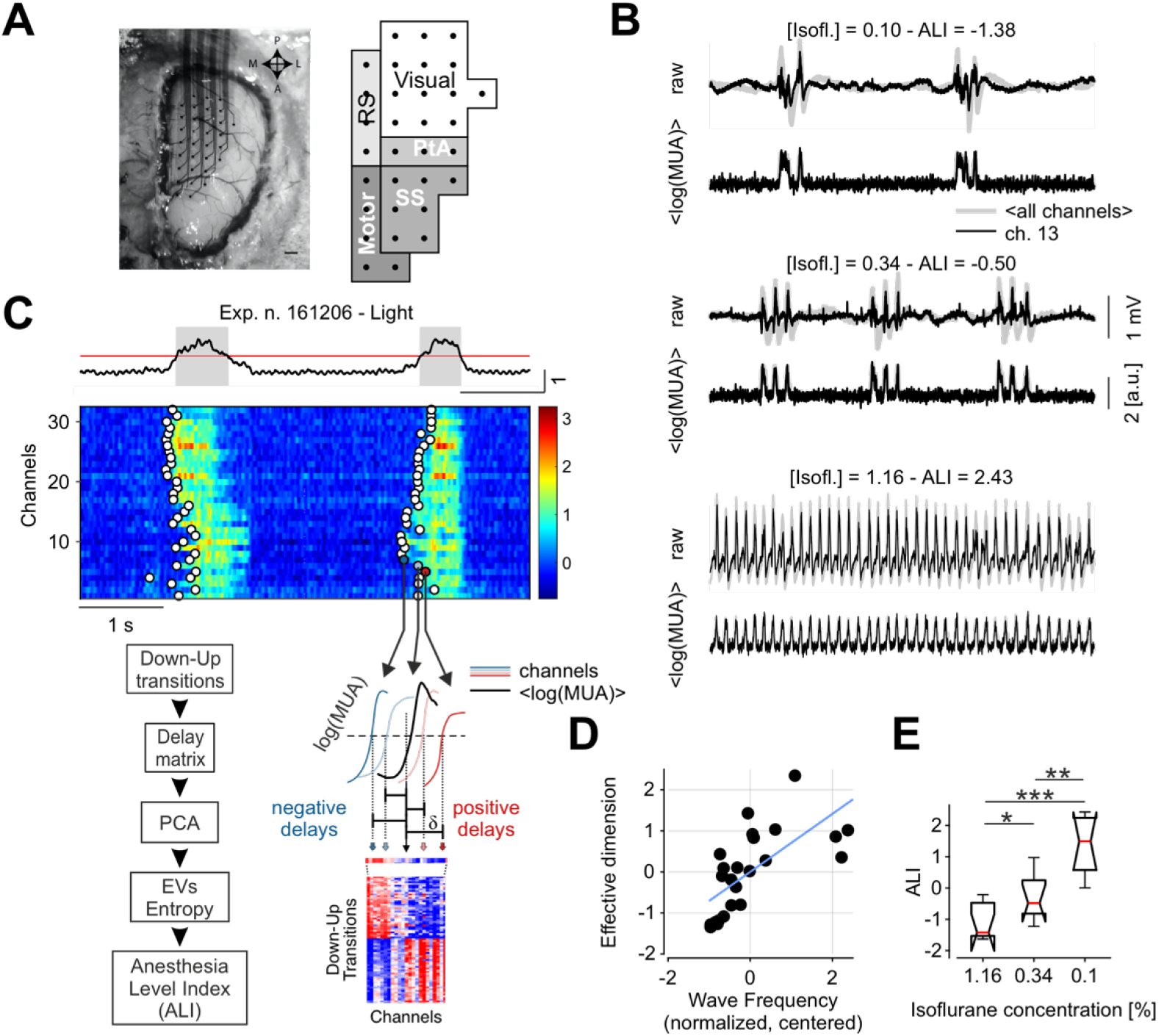
Anesthesia level index (ALI) from slow-wave dimensionality and frequency. **A.** Superficial 32-channel multi-electrode array placed on the cortical surface of anesthetized mice (left) and schematic representation of the recorded cortical areas (right, as in (Dasilva et al., 2021)). **B.** Representative raw recordings and log(MUA) for one experiment in three anesthesia levels. Grey: average of all channels; black: representative channel. **C.** Top: representative average log(MUA) (black), threshold used to extract the Up states (red) and identified Up and Down states (grey and white periods, respectively). Middle: single-channel log(MUA) of activation fronts (Down-Up transitions) from an example slow-wave (circles). Bottom right: time-lags δ of each state transition from the center of a wave compose the rows of the blue-red time-lag matrix (TLM, bottom). Bottom left: flow chart of the method used to work out the anesthesia level index (ALI) based on measure of the effective dimension of the TLM and the frequency of Down-Up waves occurrence. EV: eigenvalues **D.** Effective dimension (see Materials and Methods) and mean frequency of the Down-Up transitions normalized by their variance and centered around their mean for each recording (*n* = 24, 8 mice and 3 anesthesia levels each). Blue line: first principal component, onto which the single recordings are projected thus defining the anesthesia level index (ALI). **E.** ALI values *versus* anesthesia concentration.

As expected, both the number of effective dimensions and the frequency of Down-Up transitions fit well with the ALI (Fig. 2A, B). Furthermore, both the coefficient of variation of the Up-Down cycles (CV) and the duration of the Up states have a significant linear correlation with the ALI (Fig. 2C, D). The increase of SWA frequency is mainly due to the modulation of the Down state duration, which shows the largest exponential-like reduction with the ALI (Fig. 2E). The maximal firing rate of the cortical assemblies, measured as the peak amplitude of the Up states, shows a significant decrease as a function of the ALI (Fig. 2F).

**Figure 2:**
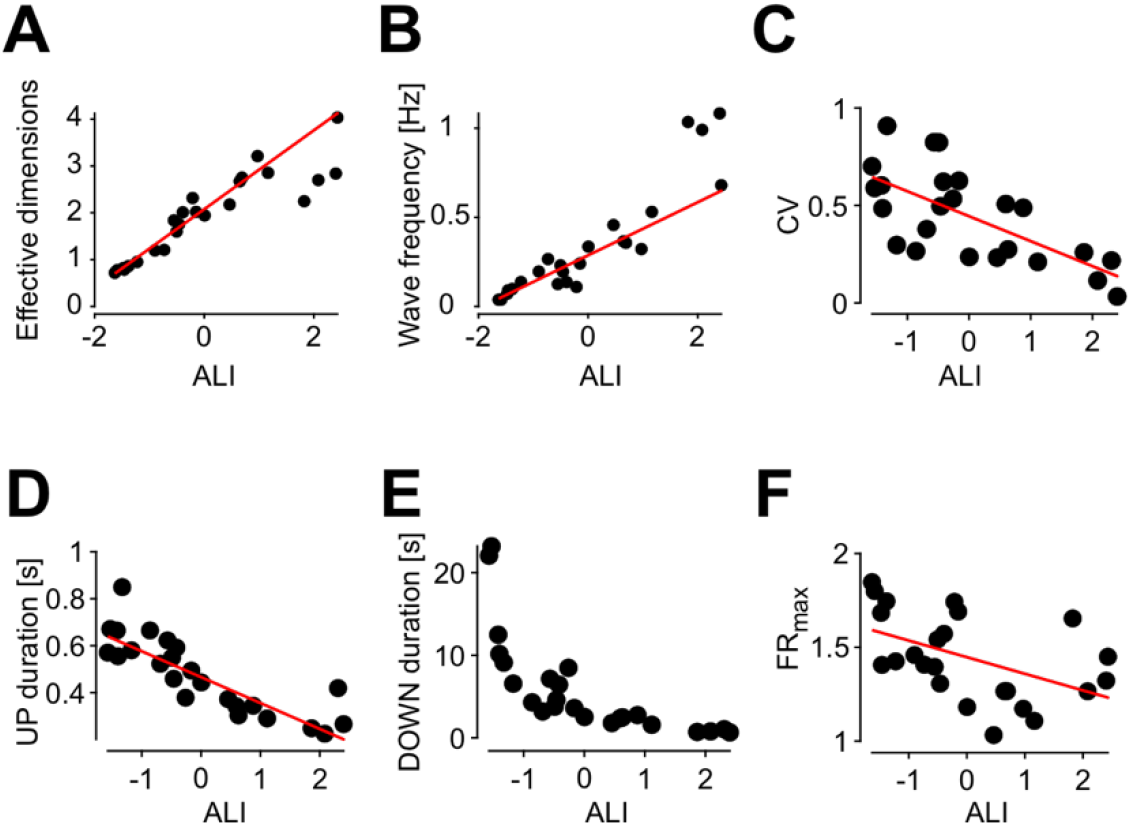
Statistical properties of the SWA as a function of the ALI. **A-F.** Number of effective dimensions (A), wave frequency (Down-Up transitions) (B), coefficient of variation of Down-Up cycles (C), mean Up (D), Down (E) state durations and maximal firing rate during the Ups *versus* the ALI (F). Red lines: linear regression (*P* < 10^−9^, *P* < 10^−9^, *P* < 0.001, *P* < 10^−6^, *P* < 0.05 for panels A-D, F, respectively).

These results highlight an acceleration and a regularization in time of the slow rhythm associated to the SWA with the lightening up of the anesthesia, similarly to what observed at the single-column level in cortex under ketamine-medetomidine anesthesia in rats (Tort-Colet et al., 2019). The awakening is accompanied by a rise of a form of complexity of the spontaneously occurring wavefronts, here defined as the number of dimensions necessary to embed them.

### Modes of propagation increase in number and speed as anesthesia fades out

Regularity in time and increase in wavefront complexity could appear as contradicting features of the slow waves. Therefore, we took a closer look at the changes occurring to the spatiotemporal patterns of neuronal activity across anesthesia levels. We did this by making use of the simultaneous acquisition of the MUA of local cortical assemblies, and by projecting their relative time of Down-Up transition into a single, common multidimensional space.

Specifically, we projected the individual Down-Up wavefronts onto the space of the principal components extracted after concatenating all TLMs (Dasilva et al., 2021), and pooling the set of recordings (*n* = 24) in three equally populated groups, based on their ALI (Fig. 3A). For the group with smallest ALI (ALI ranking 1 to 8, corresponding to the deepest anesthesia levels), we found a bimodal distribution of propagation modes in the (PC_1_, PC_2_) plane, highlighting the existence of two preferred spatiotemporal patterns. In the second group, having a higher ALI (ranking 9-16), the distribution of waves slightly moves and increases in size, while in the last group of recordings (ALI ranking 17-24) the distribution became smaller and unimodal further approaching the origin of the (PC_1_, PC_2_) plane. By visualizing the points in a three dimensional representation that included the ALI, such changes become even more apparent, as a clear progression can be observed from two “legs” (small ALI, two modes), through a fat “belly” (intermediate ALI) to a small “head” (high ALI, Fig. 3B).

**Figure 3:**
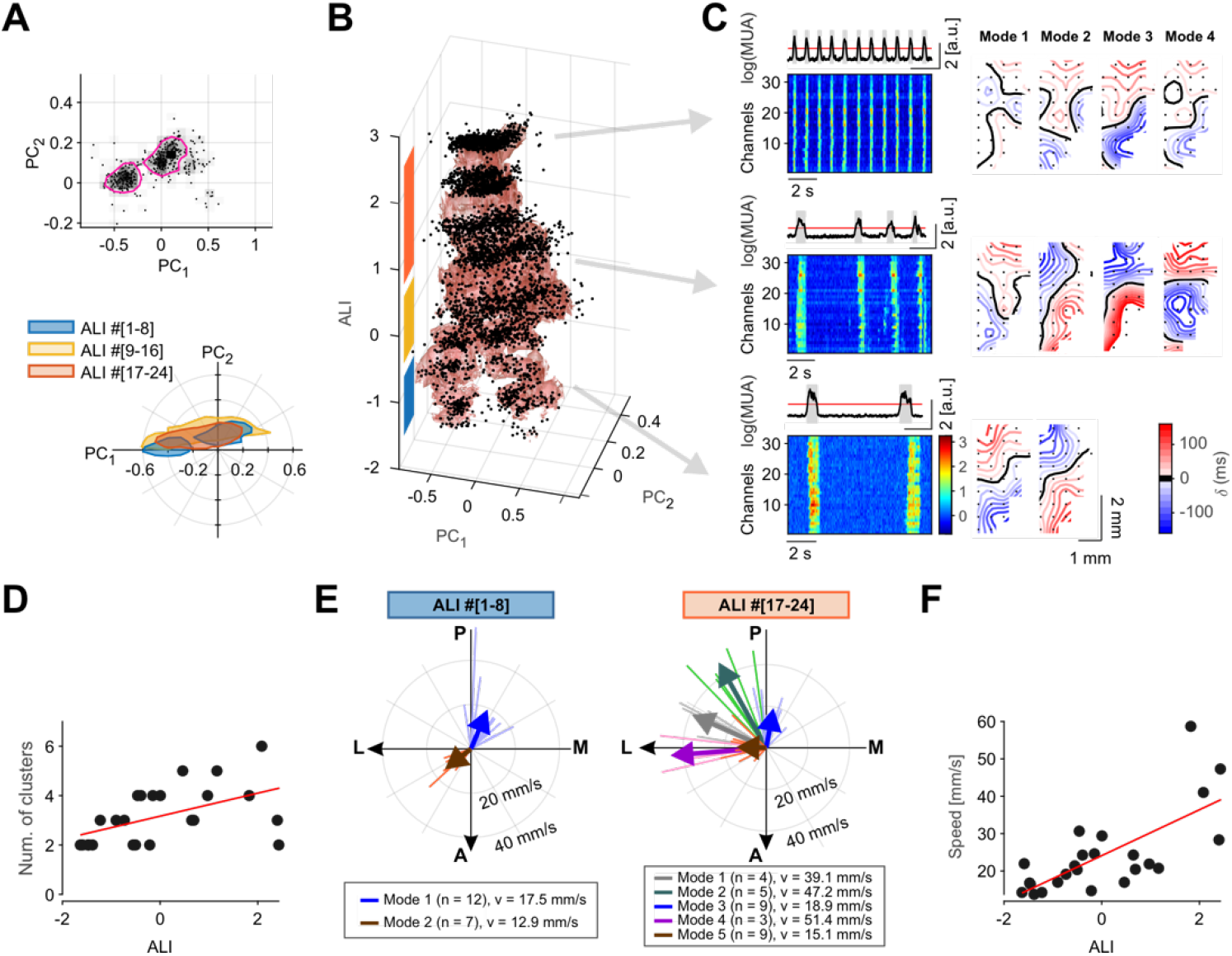
Speed and wave diversity increases as anesthesia fades out. **A.** Top: representative distribution of time-lag arrays of activation waves in the space of the first two principal components (PC_1_, PC_2_). All transitions from experiments with ALI ranking 1 to 8 are shown. Magenta contours represent 65% isodensity levels of the distribution. Bottom: 65% isodensity contours for the activation waves from experiments and anesthesia levels pooled in three equally-sized groups according to the ALI ranking. **B.** Activation waves (dots) for all experiments projected on the first 2 PCs *versus* ALI. Red shading: smoothed 65% isodensity surface. **C.** Average and single-channel MUA for three experiments and anesthesia levels representative of the ALI groups in (A) and (B). Right: resulting average spatiotemporal propagating patterns of activation waves clustered in different “modes of propagation”. **D.** Number of propagation modes per experiment and anesthesia level. Red line: linear regression (*P* < 0.05). **E.** Direction and velocity of the modes of propagation from the experiments 8 lowest and highest ALI. Modes are colored according to the result of a k means clustering. Mode clusters with less than three patterns are not shown (excluded modes: n = 1, excluded patterns: n=2). Thick arrows: average directions and velocities of each mode cluster. **F.** Mean propagation velocity for each experiment and anesthesia level. Red line: linear regression (*P* < 10^−4^).

Although the contraction of the distributions at higher ALIs might suggest the waves to be more similar to each other, this is not the case as it is apparent from the representative experiment shown in Fig. 3C, where for the three recordings at different anesthesia levels, similar patterns of propagation are grouped together resorting to automatic clustering (see Material and Methods). Indeed, the number of propagation modes, i.e. the clusters of activation wavefronts (Fig. 3C-right), increases with ALI together with the slow-wave frequency (see MUA rasters, Fig. 3C-left): a correlation confirmed also at the population level (Fig. 3D, *R*^2^ = 0.26, *P* = 0.01). This seemingly contradiction between wave distribution and number of clusters is explained by the fact that at high ALI the effective dimension is greater than two and the plane (PC_1_, PC_2_) is no more adequate to represent the variability of the activation wavefronts.

We found also a similar ALI-dependent segregation and coalescence of the directions of singled out propagation modes. By computing the average wavefront for each cluster of waves (i.e., propagation mode) and plotting the resulting average vector speeds (Fig. 3E), a prevalence of anterior-to-posterior and posterior-to-antero/lateral propagation was apparent for deep anesthesia levels (small ALI, Fig. 3E-left), as previously reported in (Greenberg et al., 2018). For lighter anesthesia levels, several medial to lateral modes appeared (Fig. 3E-right), with rare slow waves with caudal or lateral origin. Importantly, slow waves displayed a mode-independent increase of the speed with the ALI (Fig. 3F, *R*^2^ = 0.51, *P* < 10^−4^). Faster waves implies that relative delays in the TLM become progressively smaller, which in turn leads to relatively small PC_1_ and PC_2_, partly explaining why in Fig. 3B the wave distributions shrink close to the plane origin.

These results show that the rise of wavefront complexity occurring as the anesthesia fades out is due to a widening of the repertoire of the propagation modes of the SWA. Intriguingly, as wakefulness is approached such diversity is constrained to have slow waves with antero-medial origin, similarly to what found in humans under slow-wave (NREM) sleep (Massimini et al., 2004; Nir et al., 2011). In the time domain, a kind of temporal compression takes place making the slow waves more frequent and faster with the lightening up of the anesthesia.

### Wavefronts are shaped by previous waves differently across anesthesia levels

We have shown that slow waves are faster and occur closer in time during the lightening of anesthesia. Under this condition, the transit of slow waves across the cortical surface may leave a trace into the ongoing neuronal activity. This in turn could modulate the degree of wavefront complexity, thus linking together the spatial and temporal domain. To investigate such a possibility, we quantified the memory associated with the sequence of propagation modes for the different anesthesia levels.

We measured it by calculating the similarity of subsequent activation waves as the detrended correlation between the associated time-lag arrays (see Materials and Methods for details). Surprisingly, the largest amount of (negative) memory was found for deep anesthesia levels (Fig. 4A). For decreasing levels of anesthesia, the influence from the shape of previous waves firstly vanished, eventually increasing to positive memory values giving rise to a significant correlation with the ALI. A negative memory can be explained by a non-random alternation between the two modes of propagation spontaneously expressed under deep anesthesia (Fig. 4B-top). In contrast, for lighter anesthesia a propagation mode was more frequently followed by a similar one, thus leading to positive memory (Fig. 4B-bottom).

**Figure 4:**
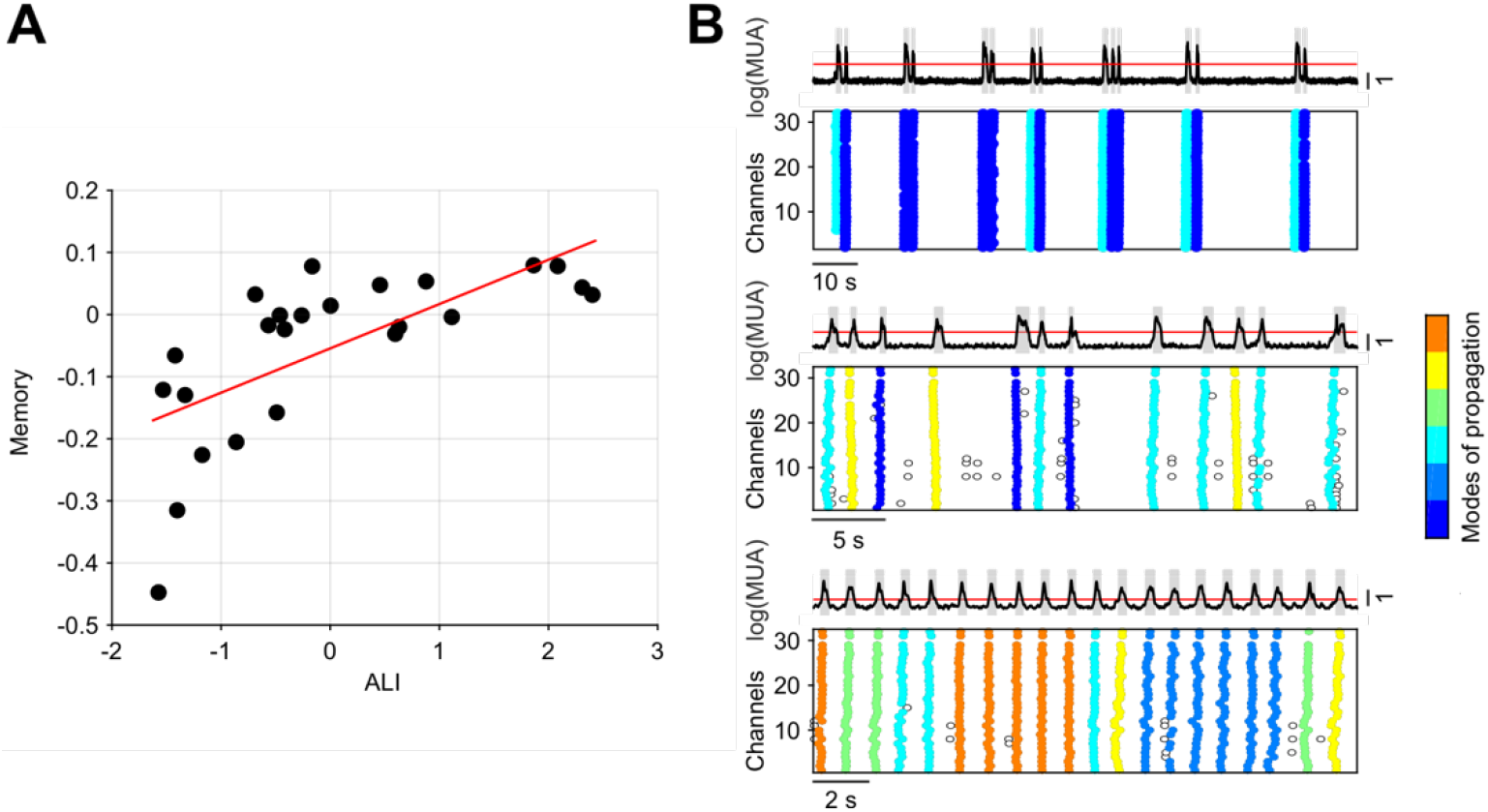
Propagation modes alternate as anesthesia level gets deeper. **A.** Autocorrelation (i.e., memory) of time-lag arrays associated to consecutive activation waves for experiment and anesthesia levels with increasing ALI. Red line: linear regression (*P* < 0.001). **B.** Average and single-channel log(MUA) for three representative experiments, showing (from top to bottom) negative, null, and positive memory. Colors represent different modes of propagations as in Fig. 3.

These results suggest that past events impact differently the onset of the current activation wave depending on the state the global network is in. The persistence and the alternation of propagation modes under light and deep anesthesia, respectively, are likely due to different mechanisms at the network and single-neuron levels. Interestingly, the former and the latter are associated to a more regular and irregular SWA rhythm, respectively, as recently pointed out in cortical slices (Capone et al., 2019).

### Leading and following assemblies differentially contribute to the rise of complexity

Having observed that the lightening of anesthesia leads to the emergence of a greater variability of the paths followed by the slow waves, we wanted to investigate whether this might be due to changes in the exogenous drive from other cortical and/or subcortical areas. Indeed, it has been previously shown that the transition to wakefulness is accompanied by a progressive restoration of the functional integration between cortical and subcortical areas (Alkire et al., 2008; Bettinardi et al., 2015; Hudetz, 2012). If this was the case, the local cell assemblies probed by the MEA would display activation and silencing MUA profiles independent from the singled out propagation modes.

To test this hypothesis, we characterized the Down-Up and Up-Down activity transitions of both the channels initiating and the channels being recruited at last by a travelling wave (referred to as leading and following channels, respectively). We observed that although there was a consistent temporal delay between leading and following channels at the Down-Up transition (activation wavefront), all channels ended the Up state simultaneously (Fig. 5A) (Sheroziya and Timofeev, 2014; Volgushev et al., 2006). In agreement with our previous results, the average amount of time the neuronal populations fired action potentials increased with the lightening of the anesthesia (due to more frequent Ups), revealing an increasing excitability of the network (Dasilva et al., 2021). However, the maximum MUA amplitude of the Ups decreased with the lightening up of the anesthesia (Fig. 5B, see also Fig. 2F). Furthermore, leading channels systematically displayed a maximum MUA significantly larger than the following ones (Fig. 5C-D, Wilcoxon rank-sum test, *P* < 10^−4^). Note that leaders and followers change from wave to wave, thus proving that the probed network contributed actively to shape the modes of propagation of the slow waves.

**Figure 5:**
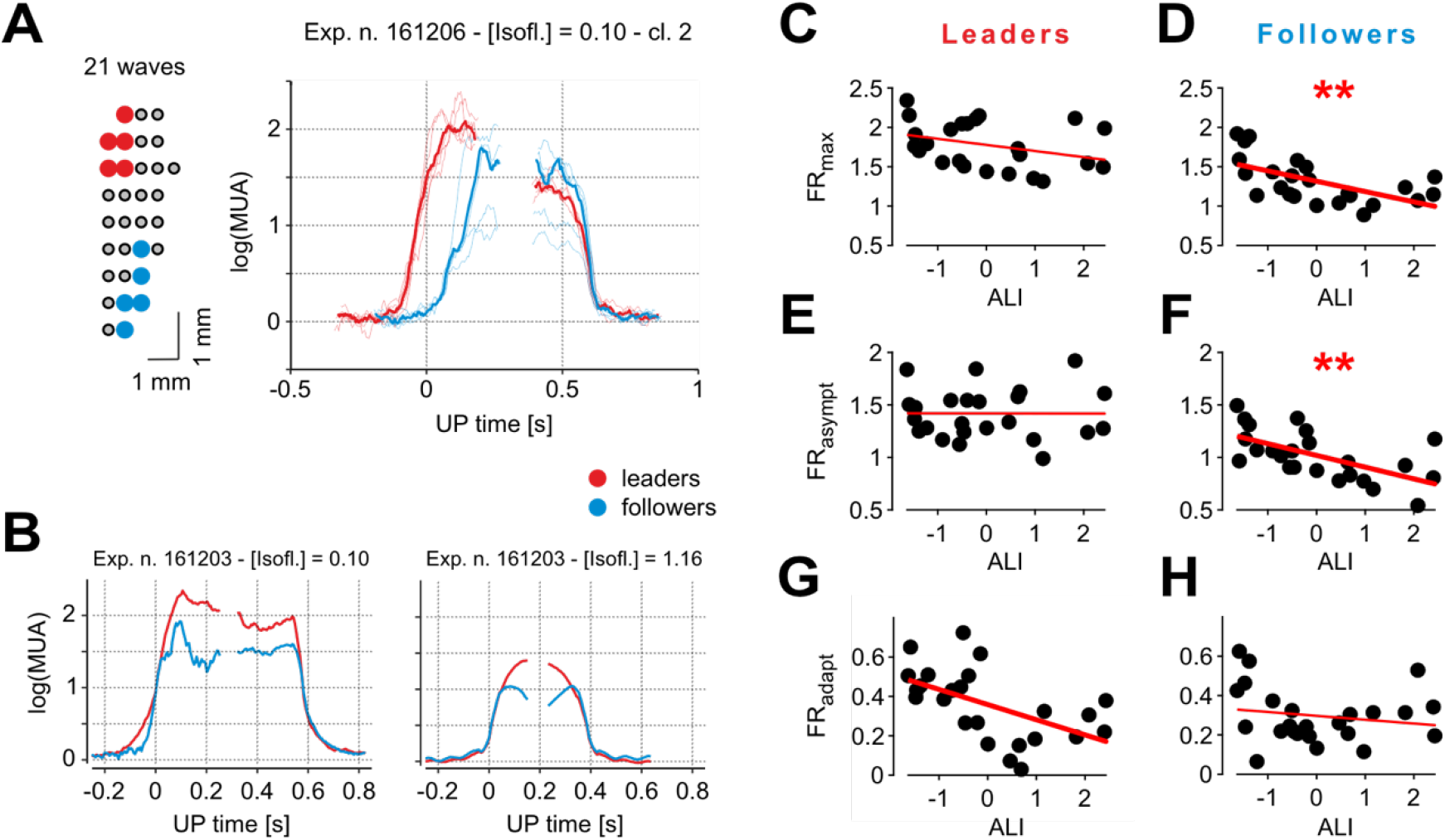
Neuronal assemblies leading wave propagations are the most excitable. **A.** Left: mapping of the five channels leading (red) and following (blue) a representative mode of propagation (in this case going posterior-anterior, and containing 21 activation waves). Right: average log(MUA) of the leading and following channels in their Down-Up and Up-Down transitions (thin lines), and their grand-averages (thick lines). The average time of Down-Up transitions in a wave is considered as time origin. **B.** Representative average log(MUA) during Down-Up and Up-Down transitions of leading and following channels for two anesthesia levels in the same animal. Differently from panel (A), time origin refers to state transitions for each group of channels. ALI values are −1.63 and 0.46 for deep and light anesthesia levels, respectively. **C-F.** Maximum (C, D) and asymptotic (E, F) firing rate (FR_max_ and FR_asympt_, respectively) during Up states for leading and following channels across all experiments and anesthesia levels (n = 24). **G, H.** Firing rate adaptation measured as the difference FR_asympt_ - FR_max_ from panels (C-F) for leading (G) and following (H) channels.

Surprisingly, it turned out that the firing rate of leading channels stays constant at all anesthesia levels (Fig. 5C, E, *P* = 0.11 and *P* = 0.99, respectively), despite the fact the Down periods progressively shrink, hence reducing the recovery time of the spiking resources responsible for the spike-frequency adaptation, i.e., determining in turn a net increase of the mean adaptation-related hyperpolarizing potassium currents. In contrast, following channels featured a strong decrease of both the maximal and the asymptotic firing rate with the ALI (Fig. 5D and F, *R*^2^ = 0.35 and 0.37, respectively, and *P* < 0.01). Consistently to these observations, the adaptation leading to a reduction of the firing rate from its maximum to its asymptotic value, is stronger for leading than for following channels (Fig. 5G and H, *R*^2^ = 0.30 and 0.03, *P* < 0.01 and *P* = 0.4, respectively).

These results suggest that as anesthesia fades out, the excitability of local cortical assemblies is diversely affected depending on the role they have in contributing to the activation wavefronts. In this way, SWA exploits the maximally available resources in terms of state-dependent excitability to find its preferential pathways of propagation.

### Changes in local excitability explain the increase of slow-wave complexity

In order to further investigate the question whether the spatiotemporal patterns observed during SWA are initiated by inputs from other (possibly subcortical) brain structures and shed light on the mechanisms underlying the initiation of the slow waves in our system, we modulated the excitability in a relatively simple network model capable to retain the relevant characteristics of the studied cortical structures. Indeed, if the observed excitability modulation of local cell assemblies was not responsible of initiating the increasingly variable propagating patterns as anesthesia is lightened, we would predict that the network would need additional external input to produce these patterns, and not a mere change in local excitability.

The *in silico* network aimed to model a wide region of the cortical surface as a 13 × 13-grid of local cell assemblies composed of 500 excitatory and 125 inhibitory randomly connected integrate-and-fire neurons (see Materials and Methods). Each of these cell assemblies modelled the cortical columns contributing to the MUA we assumed to have been recorded by a single channel of the MEA (Fig. 6A-top). We analyzed the spatiotemporal distribution of the firing activity of a 5 × 5-subset of the grid representing the cortical area probed by a schematized MEA (Bazhenov et al., 2008). We varied the working point of the model network by changing two key parameters of the local assemblies modelling the anesthesia fading out (Destexhe, 2009; Tort-Colet et al., 2019): i) the intensity of the excitatory synaptic current received from external (non-modelled) brain regions, and modulated by changing the firing rate *v_ext_* of Poisson-like upstream neurons; and ii) the strength *g_a_* of the spike-frequency adaptation due to after-hyperpolarizing potassium currents (Fig. 6A-middle, Materials and Methods).

**Figure 6:**
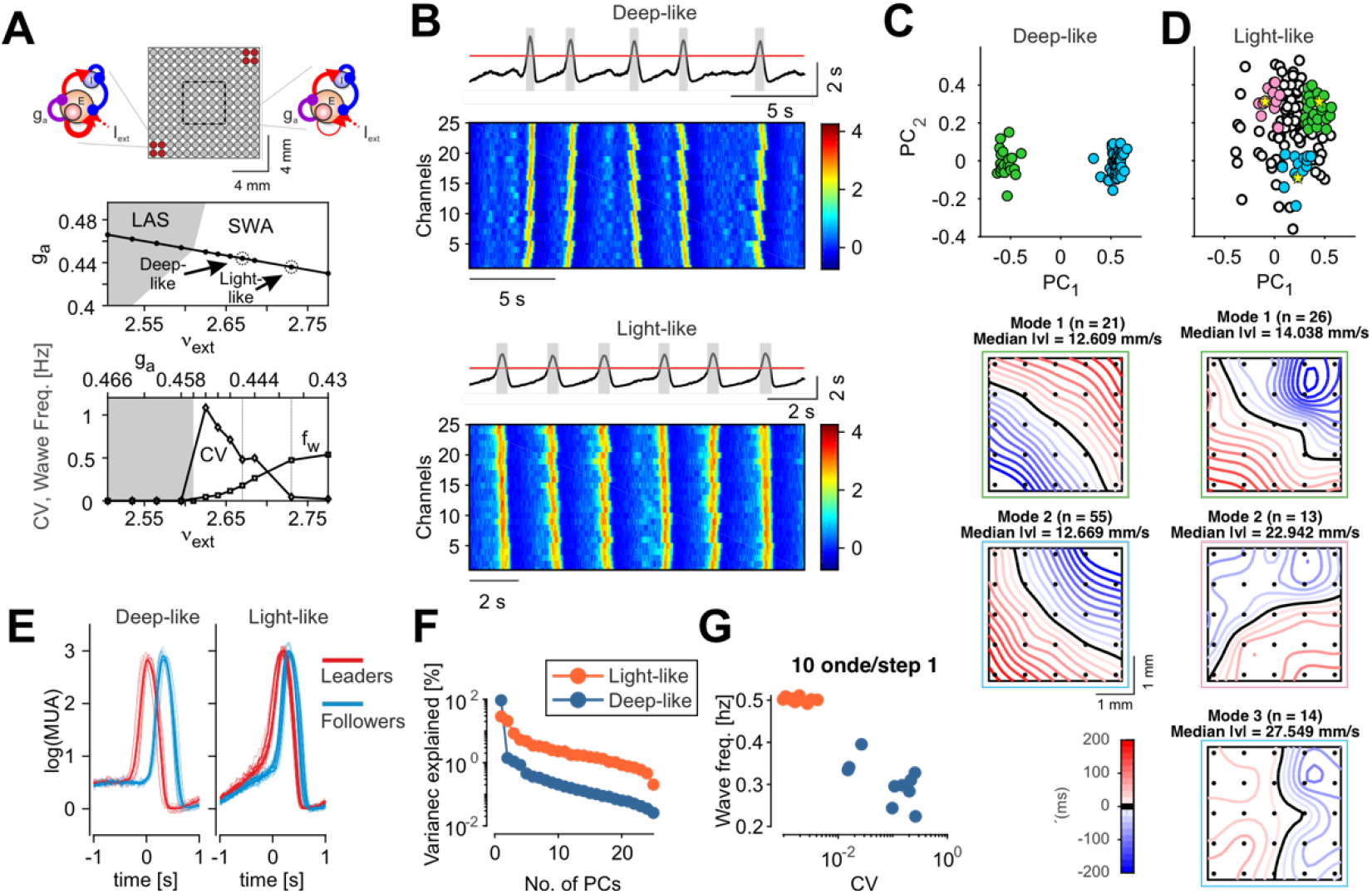
Increased excitability in model explains observed changes as anesthesia lightens. **A.** Top: schematic representation of the simulated network, consisting of 13 × 13 local cell assemblies. Cell assemblies are composed of excitatory and inhibitory integrate-and-fire neurons and are sketched on the left those with increased excitability (stronger synaptic self-excitation, red circles) and on the right the ones with reference excitability in the rest of the grid. The internal square shows the portion of the network used for the analysis. Middle: bifurcation diagram showing the different activity regimes displayed by spiking neuron network simulations as the level *g_a_* of the firing rate adaptation and the rate *v_ext_* of incoming excitatory spikes from other (external) areas are changed. To model anesthesia fading, these parameters are changed according to the depicted black line connecting the low-firing asynchronous (LAS) and the slow-wave activity (SWA) state. The arrows point to the parameter combinations used in the rest of the figure. Bottom: coefficient of variation of Up-Down cycles (circles) and frequency of waves (squares) measured in simulations along the above black trajectory. **B.** Representative average and single-channel log(MUA) in the simulated network for low wave frequency/high CV (top, “deep-like”), and high frequency/low CV (bottom, “light-like”) modelling deep and light levels of anesthesia, respectively. **C, D.** Distributions of time-lag arrays of spontaneously occurring activation waves in the model network in the plane (PC_1_, PC_2_) as in Fig. 3A-B for *in vivo* recordings. Model networks in both the deep- and the light-like conditions are shown (panel C and D, respectively). Colored dots highlight the wavefronts belonging to the modes of propagations singled out in bottom panels, relying on *k*-means clustering as in Fig. 3C. For the “light-like” case (panel D) three representative groups of wavefronts were selected; these included the points centered at the yellow stars (see Materials and Methods for details). **E.** Average log(MUA) of the five leading (red) and five following (blue) channels (thin) and averages (thick) around the Down-Up transitions for all modes shown in panel C, for the “deep-like” (left) and “light-like” (right) conditions. **F.** Percentage of explained variance as a function of the number of PCs for the TLMs extracted from the simulated data. **G.** Frequency of the activation waves *versus* the CV of the Up-Down cycles for the simulated data obtained from 10 equally-sized non-overlapping intervals of time.

In the plane (*v_ext_*, *g_a_*) the network followed a linear trajectory moving its working point from the low-firing asynchronous (LAS) region reminiscent of a “burst suppression” global state, to the slow-wave activity (SWA) regime. In this way, the lightening of the anesthesia is modelled as a simultaneous reduction of the strength of the activity-dependent adaptation *g_a_* and an increase of the external excitation *v_ext_*, which in turn can be proven to be equivalent to causing an unbalance between synaptic excitation and inhibition (Levenstein et al., 2019). This increase of local excitability in the SWA regime led to a rise of the wave frequency *f_w_* occurring more regularly in time, thus lowering the coefficient of variation CV of Up-Down slow oscillations (Fig. 6A-bottom) (Sancristóbal et al., 2016). We selected two points (referred to as deep- and light-like) on this trajectory, where the resulting CV and *f_w_* matched the experimental values (arrows on panel 6A, middle). The firing rates of the cell assemblies converted in log(*MUA*) (see Materials and Methods for details) showed spatiotemporal patterns closely resembling the experimental data (Fig. 6B). The internal variability of the patterns (i.e., the wavefront complexity) was lower under the “deep-like” condition with high CV and low *f_w_*, as demonstrated by the presence of two narrow clusters in the wave projections onto the (PC_1_, PC_2_) plane (Fig. 6C). These two propagation modes (Fig. 6C-bottom) originated by the more excitable opposite corners of the 13 × 13-grid, as this was the only structural element of heterogeneity implemented in the cortical field model.

In contrast, in the low CV/high *f_w_* (“light-like”) case, wavefront complexity was higher, as demonstrated by the wider distribution of waves in the (PC_1_, PC_2_) plane (Fig. 6D). In this state, activation wavefronts had a rather homogenous distribution of propagation directions (Fig. 6D-bottom), highlighting the possibility to have the onset of a wave from any point of the grid. The average MUA size of the Up states was comparable for the deep- and the light-like conditions (Fig. 6E). Importantly, both the decay of variance explained by the principal components (related to the effective dimensions of the wavefront space), and the dynamics in the CV-frequency plane were good markers to differentiate the two dynamical regimes (Fig. 6F, G, respectively).

These results show that the modelled cortical field was capable to reproduce the large majority of the changes observed in the SWA as anesthesia fades out. This was obtained by simply modulating the local excitability of the cell assemblies composing the network. No changes in the cortico-cortical connectivity seem to be requested. Instead, some structural heterogeneity of the local excitability needs to be incorporated as it played a role under relatively deep levels of anesthesia. Indeed, our model highlights the necessity to have a structural modulation along the rostro-caudal and medio-lateral axis (the axis connecting the two excitable corner of 13 × 13-grid) in order to obtain two preferred modes of wave propagation. Intriguingly, this is not enough to reproduce the anti-correlation of consecutive wavefronts observed in Fig. 4A at negative ALIs (memory values: 0.16 and 0.72 for light-like and deep-like, respectively). An additional component would have to be included, that takes into account active interactions with other subcortical brain structures (Adamantidis et al., 2019).

### Anesthesia fading out induces increasing fluctuations of SWA features

Our simulations confirmed the presence of a multiplicity of patterns when the excitability is bigger, i.e. in the state corresponding to light anesthesia. Together with the associated positive values of the memory (Fig. 4) this could underlie fluctuations in time of the complexity of the associated dynamical state. Think of sampling a short time segment from a pot of many possible spatiotemporal patterns with some memory and then repeating the procedure for the length of the recording. This process could result in either picking only a subset of patterns –with little variability; or picking many different ones. In the first case the embedding space (and thus, the ALI) would be small, while in the second case it would be high. Repeating the procedure, we would therefore expect a high variability of the ALI in time.

To investigate whether our data are in such a dynamical condition, we analyzed the ALI values at a finer-grained temporal scale along the whole time of the recordings. This was done by computing the effective dimension and the slow-wave frequency in sliding windows containing a fixed number of waves (Fig. 7A). Only a few recordings (3 out of 24) showed a marked non-stationarity, highlighted by the isolated trajectories in the frequency-effective dimension plane starting in one point but ending up somewhere else on the plane (see also an example of trace in Fig. 7A, inset). When analyzing the ALI values resolved in time for the rest of the recordings, these segregated in two groups, pretty much separating those with the ALI<1 from those with the ALI>1 (Fig. 7B). This effect was quantified by fitting the ALI distribution for the first 200 s versus the last 200 s of the acquisition time with a mixture of up to four Gaussian distributions (Fig. 7C, see Materials and Methods for details). Additionally, the recordings having average ALI above 1 tended to further increase their ALI at later times (Fig. 7B). Critically, the size of the fluctuations around the fitted trend of the ALIs significantly increased with the ALI itself (Fig. 7D, *R*^2^ = 0.49, *P* = 0.001). Importantly, when computing the ALI value resolved in time for our network simulations, these nicely fitted in the high- and low-value groups (Fig. 7B, green traces). Furthermore, the fluctuations of the simulated data also showed a clear increase for the “light-like” case (Fig. 7D, green crosses).

**Figure 7:**
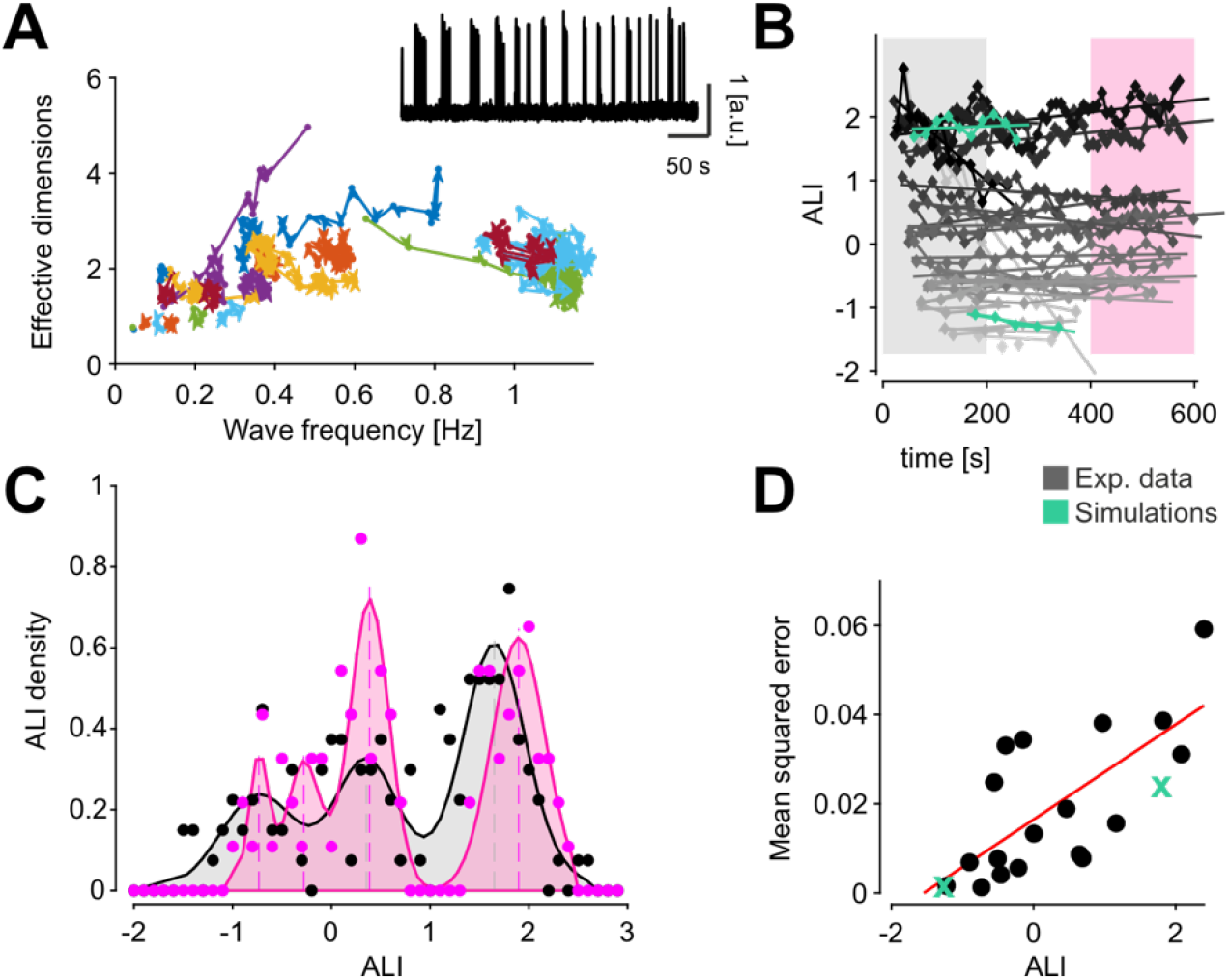
ALI drifts in time towards two preferred values at fixed anesthesia levels. **A.** Time-resolved relationship between effective dimension of activation waves and their frequency of occurrence for all experiments and anesthesia levels (n = 24) each identified by a different color. Points of the trajectories are computed from time windows including 30 consecutives activation waves. Adjacent time windows have 10 waves of overlap. Inset: example experiment with log(MUA) displaying non-stationary frequency of wave occurrence. **B.** Time-resolved ALI computed in the same time windows as in (A). Time values are the centers of the time windows. Lines are linear fits for each experiment and anesthesia level. Colored intervals highlight the first and the last 200 s of the experiments. Grey levels represent the average ALI value. Green: simulations **C.** Distribution of ALI values for the first and the last 200 s of the experiments (highlighted in panel B). Colored shading, fit of the distribution with up to four Gaussian probability densities (ALI values of the peaks: for the interval [0, 200] s: −0.73, 0.35, 1.65; for the interval [400, 600] s: −0.74, −0.28, 0.38, 1.9). **D.** Mean squared error of the linear fits showed in (B) as a function of the mean ALI per experiment and anesthesia levels (n = 24). Red line: linear regression. Green: simulations.

In summary, the lightening up of the anesthesia leads to larger fluctuations of the ALI in time, compatibly with the hypothesis that the increased slow-wave complexity is due to spatio-temporal patterns of activation (i.e., states) more prone to wander across their state space, as perfectly predicted by network simulations. Intriguingly, such a dynamical condition appears to be strengthened in the recordings with ALI above the value of 1.

## Discussion

The unconscious state is characterized by slow waves of brain activity spanning a continuum spectrum of dynamical features such as their frequency of occurrence and the distribution of their propagation speed and direction (Dasilva et al., 2021). Here, we investigated how such a continuum emerges in the same network of cortical cell assemblies as the level of isoflurane anesthesia reduces, thus mimicking the sleep-wake arousal process (Torao-Angosto et al., 2021). Besides a progressive increase of the “spatial” richness of the spontaneously expressed propagation modes, measured as the dimensionality of the slow-wave features, we found an unexpected change of their sequential memory: while at deep anesthesia consecutive waves inverted their direction of propagation (front to back vs back to front), at light anesthesia consecutive waves were more similar. The rise of spatial complexity with lighter anesthesia levels, that we measured here as an increase of the effective dimension of the space embedding the activation wavefronts, might then appear to be in contradiction with the observed temporal “simplicity” associated to the fact that consecutive waves tend to be more similar to each other close to the awakening. In a previous study we argued that a repetition of similar waves can be due to adaptation effects (Capone et al., 2019). Indeed, the propagation of more frequent slow waves leads to a more fatigued cortical tissue where excitability is firstly recovered where the last wave initiated, such that the next activation will likely start from the same place. In this way, increased memory corresponds to longer time scales needed to “forget” previous events. Having longer time scales does not mean that the space embedding the activation wavefronts cannot be fully explored. It only means that more time is needed in order to express the available dynamical richness, therefore to express waves of different origin and propagation. Note that longer time scales imply stronger correlations between spatiotemporal patterns which is a typical fingerprint of the critical dynamics arising when the transition from the synchronized and the desynchronized global state is approached (di Santo et al., 2018): a possible other side of the “complexity” coin.

Our network simulations of the cell assemblies in a large cortical field closely reproduced the large majority of our experimental observations. The fact that this was obtained without modifying the underlying “structure”, i.e. the connections between the elements of the network, highlights the local excitability modulated by activity-dependent adaptation as a key element to explain the functional changes observed during the arousal from deep anesthesia. In this regard, the channels leading in time the onset of Up states, were those with the highest firing rates (Fig. 5C), similarly to what found in cortical slices where the leaders of the activation wavefront lie in layer 5, and have longer Up states (Capone et al., 2019). Excitability of leading channels is expected to be due to activity reverberation in the high firing state which can persist in time like in bistable units (Jercog et al., 2017; Levenstein et al., 2019; Mattia and Sanchez-Vives, 2012). Even if travelling waves do not originate from the cortical location probed by the multi-electrode array and hosting the wave-dependent leading channels, they likely find a preferential pathway marked by the most excitable cell assemblies.

### Switch of global brain state associated with anesthesia

The changes in the memory effects on the past slow waves can also explain the increase of variability in the ALI as the anesthesia fades out. Indeed, under light anesthesia the small bunches of consecutive slow waves considered to produce single points of Figure 7 are often composed of similar wavefronts (thus yielding positive memory values) residing into a limited region of the whole embedding space, which in turn is rich and wide. The size of this region varies from bunch to bunch resulting into a highly variable ALI in time. Differently, under deep anesthesia the slow waves are concentrated in two focused clusters, and being consecutive wavefronts anti-correlated (negative memory), the cortical network jumps from one cluster to the other. In this condition each bunch of few waves is sufficient to probe the whole embedding space, leading to small changes in the ALI from one bunch to another.

The progressive slowdown of the wandering across the wavefront space as the ALI increases, recalls the dynamics of a system in a shallowing valley of a potential energy landscape close to a critical transition (Scheffer et al., 2009). This would be compatible with the hypothesis that unconscious (sleep) and conscious (wake) states are two distinct conditions coexisting as attractors of a global brain dynamics (Adamantidis et al., 2019; Saper et al., 2010). Like in a flip-flop ‘switch’, intermediate states tend to be avoided as the global attractors are separated by a repulsive barrier. A similar bistability has been invoked to explain the hysteresis loop observed in the switch from anesthesia to wakefulness and vice versa (Friedman et al., 2010; Hudson et al., 2014; McKay et al., 2006; Proekt and Kelz, 2018). In this framework the existence of a barrier was tested via electric stimulation of the ventral tegmental area, which would induce the arousal from the unresponsive anaesthetized state only when intense enough (Joiner et al., 2013; Solt et al., 2014). Intriguingly, our finding of divergent trajectories above and below the ALI ≈ 1 shown in Figure 7 corroborates the hypothesis of a reflecting barrier between anesthesia and wakefulness, highlighting a possible destabilization of the (attractive) unresponsive state as the anesthesia level lowers below a threshold dose. Of course, further and more focused experiments are required in order to confirm such theoretical framework.

### Role of subcortical regions in the slow-wave anti-correlation

Another intriguing aspect of the correlation between consecutive wavefronts is the significant negative memory effect found under deep anesthesia. Although slow-wave activity features like mean frequency and Up (Down) durations measured in this conditions are quantitatively similar to the one we observed in isolated cortical slices (Sanchez-Vives et al., 2010; Sanchez-Vives and McCormick, 2000), we had never found an anti-correlation of consecutive slow waves *in vitro* (Capone et al., 2019). Our network model was unable to reproduce such negative memory effect in a deep-like dynamical conditions, although a suited degree of heterogeneity in the cell assembly organization was incorporated in order to have two clusters of propagation modes (see Figure 6). Thus, a question arises: where does this anti-correlation come from?

To address this issue, we first noted that under deep anesthesia the slow-wave activity displays the typical hallmarks of burst suppression with prolonged Down states alternating with periods of brief bursts of spikes and waves (Lewis et al., 2013; Purdon et al., 2015; Steriade et al., 1994). Similarly to mice, burst suppression induced in humans by propofol (Lewis et al., 2013) has suppression (Down) periods with median duration of 4.76 s, and activity bursts (Up) lasting more than half a second. Furthermore, burst onsets propagate across the cortex as an activation wave travelling roughly 1 cm in 225 ms. The resulting speed of 44 mm/s is faster than the 15 mm/s we measured, but it should be considered as an upper bound due to the folded structure of the human cortical surface – a characteristic which is almost absent in mice. In our network model the deep-like global state is reproduced by making the Down state metastable in its local cell assemblies, such that Up states are elicited by endogenous fluctuations of the spiking activity, following a self-inhibited period in which the adaptation level (related to the K+ afterhyperpolarization current, AHP) relaxes to its resting value. Compatibly with this picture, under burst suppression an exogenous stimulation elicits an activity burst only if a refractory period is passed (Kroeger and Amzica, 2007), and if its intensity is above a threshold value (Amzica, 2015).

The presence of isolated Up states (bursts) of cortical activity under deep anesthesia can in principle provide supra-threshold input to other subcortical areas. More specifically, the back-to-front waves ending in the frontal cortex can elicit a reaction in the hippocampal network. If this activity would persist long enough (i.e., beyond the refractory period, after which the full excitability is recovered) and it would be fed back to the cortex, this will ignite a new wave propagating backward. Evidence of such cortico-hippocampal interaction has been described in rodents with simultaneous recordings in the hippocampus (CA1) and in parietal (Hahn et al., 2007), temporal (entorhinal) (Hahn et al., 2012), visual (Ji and Wilson, 2007) and in the whole (Busche et al., 2015) cortex. In a similar way, other subcortical structures like first-order thalamus could be involved when fronto-caudal propagation modes occur (Sheroziya and Timofeev, 2014; Steriade et al., 1993b), giving rise to a rebound activity, which is then back-projected onto cortical neurons (Steriade et al., 1993a).

If the suggested scenario was confirmed, negative memory of consecutive slow waves under burst suppression would be the results of a state-dependent chat between cortical and subcortical structures. This phenomenon should leave a recognizable fingerprint: activity bursts would occur often as duplets interspersed by relatively long Down (suppression) periods as in the example recording shown in Figure 1B-top.

### Relationship between slow waves and EEG microstates

As cortical Down-Up transitions during slow-wave activity are known to induce thalamic-generated spindle oscillations (7-16 Hz) (Andrillon et al., 2011; Mak-Mccully et al., 2017; Muller et al., 2016), in view of our current results, a possible relationship may be seen between spatiotemporal phase patterns of these two rhythms. Indeed in ECoG recordings during quiet wakefulness in humans and macaques, alpha oscillations (7-13 Hz) display a preferential direction of propagation from antero-superior to postero-inferior cortex (Halgren et al., 2019). This distribution of directions widely overlaps with the one we found for slow waves under relatively high ALI, i.e., under light anesthesia, compatibly with the evidence that quiet wakefulness is a kind of nonstationary wandering between first sleep stages and wakefulness (Reimer et al., 2014; Vyazovskiy et al., 2011). One issue here is that reported alpha waves have average speed of propagation of about 0.9 m/s (Halgren et al., 2019), far faster than the speed of the activation wavefronts we measured, ranging on average from 15 to 40 mm/s.

A second analogy comes from the field of EEG electrophysiology, where alpha oscillations are associated to the so-called EEG microstates, i.e., spatial snapshots of the field potential phases captured at the peaks of these waves, eventually clustered in a few number of stereotyped spatiotemporal patterns (Brodbeck et al., 2012; Lehmann et al., 1987; von Wegner et al., 2017). Here, the spontaneous brain activity (i.e., sleep and quiet wakefulness) is hypothesized to be a continuous transition between such EEG microstates (Brodbeck et al., 2012; Lehmann et al., 1987). Intriguingly, EEG microstates display some degree of memory about past events, analogously to what we report here for cortical slow waves. Furthermore, EEG microstates are non-Markovian (von Wegner et al., 2017) with each microstate likely persistently reoccurring for about half a second: a time scale remarkably reminiscent of the Up state duration measured during SWA. Also, this timescale is not independent from the brain state: EEG microstates persist for longer times in deep sleep (stage N3) compared to wakefulness and N1 stage (Brodbeck et al., 2012). This in turn is reminiscent of the elongation of the Up state duration we found as the ALI decreases from light to deep anesthesia.

In summary, we propose that slow waves can be seen as “mesostates”, probably embedding coherent spindle-related EEG microstates, which coarse-grain the richness of propagation modes we observed in invasive recordings. The fine-grained resolution characterizing the slow waves allows us to generalize the idea that the rise of complexity associated to the transition from the unconscious to conscious brain state is due to a continuous widening and reshaping of the landscape of such mesostates, which by the way can be fully described by a parametric modulation of the local excitability of cortical assemblies.

## Materials and Methods

### Animals

All procedures were approved by the Ethics Committee at the Hospital Clinic of Barcelona and were carried out to the standards laid down in Spanish regulatory laws (BOE 34/11370-421, 2013) and in the European Union directive 2010/63/EU. Recordings were performed in eight adult male C57BL/6J mice bred in-house at the University of Barcelona and kept under 12h light/dark cycle with food and water ad libitum. All experiments were performed in the animal’s subjective daytime.

### Surgical procedures

Anesthesia was induced with an intraperitoneal injection of ketamine (75 mg/kg) and medetomidine (1.3 mg/kg) and maintained by the inhalation of isoflurane in pure oxygen. Atropine (0.3 mg/kg), methylprednisolone (30 mg/kg) and mannitol (0.5 g/kg) were administered subcutaneously to avoid respiratory secretions and edema. Body temperature was constantly monitored and kept at 37 °C using a thermal blanket (RWD Life Science, China). Once under the surgical plane of anesthesia, mice were placed on a stereotaxic frame (SR-6M, Narishige, Japan) and a craniotomy and durotomy were performed over either the right or left hemisphere from −3.0 mm to +3.0 mm relative to bregma and +3.0 mm relative to midline. This broad craniotomy and durotomy were chosen to allow access to a large area of the targeted hemisphere, either left or right. Three levels of anesthesia were reached that were classified according to the provided isoflurane concentrations: Deep = 1.16 ± 0.08% (mean ± s.e.m); Medium = 0.34 ± 0.06%; Light = 0.1–0% (Dasilva et al., 2021). The volume delivered was 0.8 l/min and the animals were breathing freely via a tracheotomy. Each anesthesia level was maintained for 20–30 minutes and recordings were each of 300–500 s duration consistently obtained in a stable slow oscillatory regime (~10 minutes after the change in concentration) and thus, before microarousals appeared due to lighter anesthesia (Tort-Colet et al., 2019). Furthermore, the absence of reflexes was regularly checked. With respect to the extreme of the depth of anesthesia, we had already determined in previous experiments the range of anesthetic that could be used to ensure the maintenance of a bistable pattern of activity, while avoiding respiratory depression.

### Electrophysiological recordings

Extracellular LFP activity was recorded by means of 32-channel multi-electrode arrays (Pazzini et al., 2018) (550 μm spacing between recording points) covering the entire exposed area of the cortex. The signal was amplified by 100 and high-pass filtered above 0.1 Hz (Multichannel Systems, GmbH), digitized at 5 kHz and fed into a computer via a digitizer interface (CED 1401 and Spike2 software, Cambridge Electronic Design, UK). We ensured that all 32 channels were working and recording good signal during the experiments.

### Data analysis and statistics

All the analyses were performed in MATLAB (The MathWorks, Natick, MA). Linear fits shown throughout the manuscript were computed using a linear regression model reporting both *R*^2^ and *P* values of the fits.

#### MUA estimate and detection of the state transitions

Up and Down states and multi-unit activity (MUA) were estimated from the recorded raw signals (Mattia et al., 2010; Reig et al., 2010; Sanchez-Vives et al., 2010). Briefly, the power spectra were computed from 5-ms sliding windows of the raw signal. MUA was estimated as the relative change of the power in the [0.2, 1.5] kHz frequency band. Such spectral estimate of the MUA was not affected by the electrode filtering properties, and it provided a good estimate of the relative firing rate of the pool of neurons nearby the electrode tip. In a second step, the MUA was logarithmically scaled to compensate for the high positive fluctuations due to the neurons closest to the electrode, thus obtaining the log(MUA) signal. We then smoothed the log(MUA) by performing a moving average with a sliding window of 40 ms. Reference value for this signal was associated to the most prominent peak of its distribution corresponding to the average activity during Down states and set to 0. From the long-tailed histogram of log(MUA), an optimal threshold separating Up and Down activity states was set at the absolute value of 0.4. The threshold for detecting both Down-to-Up and Up-to-Down transitions was the same.

The frequency of the SO was the inverse of the duration of the entire Up-Down cycle. We reconstructed the activation waves from the detected Down-to-Up state transitions as in Capone et al. (2019) and in Dasilva et al., 2021. Briefly, each wave was reconstructed pooling together the transitions occurring in multiple electrodes in a reasonable time interval that was iteratively reduced until each wave contained no more than one transition per channel. Each reconstructed wave was associated with a vector of relative time lags computed as the difference between the time of occurrence of the wave in each electrode and the average time of occurrence across all the electrodes taking part in the wave propagation. We rejected the waves occurring in less than twelve channels, otherwise we replaced the missing values with the result of a nearest-neighbor interpolation using the five nearest points in terms of Euclidian distance of wavefronts/time lag vectors. Using the resulting vectors describing the detected activation wavefronts, for each animal and anesthesia level we composed a time-lag matrix (TLM) having a number of rows equal to the number of waves and a number of columns equal to the number of recording channels of the experiment. The spatiotemporal course of the wavefronts was then obtained by spatially interpolating the time lags without smoothing, using a thin-plate spline method.

#### Extraction of the anesthesia level index (ALI)

In order to determine with more accuracy, the level of anesthesia as perceived by the animal, we introduced an index based on the data, referred to as the anesthesia level index (ALI). This was computed with the following steps: first, we computed for each dataset the fraction of variance *p_k_* = *λ_k_*/ *Σ_j_ λ_j_* explained by each principal component (*λ_k_* are the eigenvalues of the covariance matrix) of the TLM. Next, we computed the effective dimension as *D* = *e*^(*H*−1)^ where *H* = − *Σ_k_ p_k_* log*p_k_* is the Shannon entropy of the variance distribution. Under the hypothesis of an exponential distribution of the variance, *p_k_* ∝ *e*^−*k*/*D*^provided that *D* ≪ *N*, where *N* is the number of recording channels considered (*N* ≤ 32). Both these conditions where well satisfied by all recording sessions analyzed in this work, and according to (Schiff et al., 2007), *D* is the minimum number of modes explaining the 90% of the system energy, thus representing the number of dimensions needed to describe its dynamics. We then defined a 2-columns matrix *W*, whose columns contain the wave frequencies and the effective dimension *D* for each wave, both normalized by their respective standard deviations. Finally, we computed the principal components of *W* and the ALI was defined as the projection of each row/dataset of *W* onto the resulting first principal component.

#### Modes of propagation and speed of the modes

Relying on the first principal component of TLM we clustered waves/rows sharing similar features with a k-means algorithm. The optimal number of clusters to be used to group the waves in each anesthesia level was chosen using a Silhouette method. For each cluster, we obtained the mean wavefront and its spatiotemporal profile by averaging the waves belonging to it. The mean wavefront extracted from each cluster was used to compute the mean velocity and direction of propagation of the cluster as in (Capone et al., 2019). This was done relying on the smoothed surface *T*(*x, y*) (thin-plate smoothing spline) of the relative time lags associated with each cluster and distributed according to the electrode positions, and computing the local velocity as *V*(*x, y*) = 1/(*∂_x_T^2^ + ∂_y_T^2^*. The gradient of this field points out the direction of the wave propagation. Mean values are computed across the 32 electrode positions.

#### Memory of the patterns

We defined the memory of the spatiotemporal patterns as the average overlap of consecutive wavefronts, where for the latter we used the rows of the TLM defined above, corrected with a shuffling method. Specifically, for each dataset we computed the dot product of the unitary vector containing the Down-Up lag times and all subsequent lag vectors. In order to prevent large effects of noise, we replaced the rows of the TLM with the average over the nearest (in the Euclidean sense) five other rows. Next, we subtracted from the average distance to the following waves the mean of a shuffled version of the distance to each wave. The latter was computed as the average of 25 repetitions of the distance between every wave and a randomly chosen other wave in the experiment.

#### Leaders/followers analysis

For the analyses related to the leading and following cell assembly reported in Figure 5, we extracted from the TLM of each dataset the first and the last five channels participating to every wavefront and defined them as leaders and followers, respectively. In Figure 5A and B the average waveforms of the leaders and the followers for representative clusters (as defined above) of wavefronts are shown. The maximal firing rate during the Ups (Fig. 2F and Fig. 5C,D) was then computed as the maximal amplitude of the log(MUA), whereas the asymptotic firing rate (panels E, F) is the average log(MUA) amplitude in the interval [−200, −100] ms before the Up-Down transition.

### Network model and simulations

Network model shown in Figure 6 was composed of standard point-like integrated-and-fire (LIF) neurons. It is organized as a set of interacting modules, each consisting of two interconnected pools of excitatory and inhibitory spiking neurons. Similarly to (Mattia et al., 2013), each cortical module was composed of 250 inhibitory (I, 20%) and 1,000 excitatory (E, 80%) neurons with spike frequency adaptation (SFA). The neurons in the excitatory population were divided into a “foreground” (F, 25%) and “background” (B, 75%) sub-pools. Such modules were spatially distributed on a 2-dimensional grid of 13 × 13 sites. The inter-modular connectivity was greater than zero only between pairs of populations displaced at relative distance less than 3, thus modelling cortico-cortical horizontal connections. Membrane potential *V*(*t*) of each neuron evolved according to 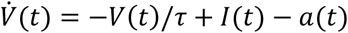 where 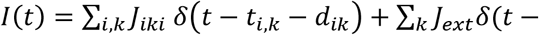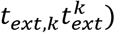 is the synaptic current from recurrent (*i*) and external (*ext*) presynaptic neurons emitting their *k*-th at time *t_i,k_* and *t_ext,k_*, and transmitted with efficacy *J_i_* and *J_ext_*, respectively. Recurrent spikes are delivered with axonal transmission delay *d_i_*. The adaptation current *τ_a_*ȧ(*t*) = −*a*(*t*) + *g_a_ τ_a_* ∑_*ik*_ (*t* − *t_ik_*) models the activity-dependent afterhyperpolarizing *K*^+^ current underlying SFA, increasing by *g_a_* = 0.04 mV/ms at each spike emission time *t_k_* of the neuron and relaxing with a characteristic time *τ_a_* = 500 ms. No SFA was incorporated in inhibitory neurons. Membrane potential decay constant was *τ* = 20 and 10 ms E and I neurons, respectively. Spikes were emitted when *V*(*t*) crossed the threshold Ɵ = 20 mV, after which potential was reset to 16 mV for an absolute refractory period *τ*_0_ = 2 and 1 ms for E and I neurons, respectively. External spikes mimicked the input from the other cerebral areas and were implemented as a Poisson process from *C_ext_* independent source firing at rate *v_ext_*. We choose *C_ext_* = 800, *v_ext_* = 3 *HZ*, *J_ext_* = 0.43 mV and 0.56 mV for E and I neurons, respectively, in order to have a fixed point with firing rate for E and I neurons at *v_E_* = 3 Hz and *v_I_* = 6 Hz, respectively, under mean-field approximation (Amit and Brunel, 1997). To this purpose the synaptic efficacies were randomly chosen from a Gaussian distribution with mean *J_αβ_* and standard deviation Δ*J_αβ_* depending on the type of presynaptic (*β* = *E*, *I*) and postsynaptic (*α* = *E*, *I*) neurons, whereas Δ*J_αβ_* = 0.25 *J_αβ_* for any **α** and *β*. Inhibitory connections were set to *J_II_* = *J_EI_* = −1.5 mV, *J_EE_* = 0.43 mV and *J_IE_* = 0.56 mV. In the same mean-field framework we structured the cortical module to have an additional high-frequency fixed point for the F excitatory neurons by potentiating synaptic self-excitation *J_FF_* and depressing the other excitatory connections *J_FF_* and *J_BF_* compared to *J_EE_* as in (Amit and Brunel, 1997): *J_FF_* = 0.61 mV and *J_FF_* = *J_BF_* = 0.39 mV. Intra-modular connectivity between excitatory neurons (i.e., the probability be synaptically coupled) was *c_EE_* = 0.8, whilst the inter-modular connectivity decreased with distance (Ercsey-Ravasz et al., 2013) assuming the values (0.03, 0.018, 0.007, 0.004, 0.001, 0.001). Axonal transmission delays were drawn from an exponential distribution with mean 〈*d_αE_*〉 = 22.6 ms and 〈*d_αI_*〉 = 5.7 ms (*α* = *E*, *I*). In the 13 × 13 grid, the modules in the two opposite 2 × 2 corners shown in Fig. 6A have an enhanced self-excitation among F neurons (*J_FF_* = 0.65 mV) to increase their excitability. Analyses were performed in the 5 × 5 subset of modules in the center of the grid. In order to use the same analytical approach developed for *in vivo* recordings, we transduced the simulated activity in an *in silico* representation of MUA. MUA was computed from the firing rates *v_f_*(*t*) of each module, subsampled at 5 *ms*, by adding a white noise *ζ*(*t*) with 〈*ζ*(*t*)〉 = 0 and 〈*ζ*^2^(*t*)〉 = *η*^2^, where *η* = 0.4 ∙ min(*v_F_*(*t*)), to mimic the fluctuations observed in the experiments.

Simulations were performed with NEST (Gewaltig and Diesmann, 2007). Full Python code where all the missing model parameters and information is freely available in EBRAINS Knowledge Graph (https://search.kg.ebrains.eu/instances/Model/eb925263fd703c9c7aec2ee94e8be72520e3b69e).

## Acknowledgments

Funded by EU H2020 Research and Innovation Programme, Grant 785907 (HBP SGA2) and 945539 (HBP SGA3) to MVSV and to MM and by TRANSLACORTEX by the Spanish Ministry of Science and Innovation through the MICINN under grant BFU2017-85048-R.

## Notes

### Competing Interest Statement

The authors have declared no competing interest.

